# Physical activity and risk of Alzheimer’s disease: a two-sample Mendelian randomization study

**DOI:** 10.1101/819821

**Authors:** Sebastian E Baumeister, André Karch, Martin Bahls, Alexander Teumer, Michael F Leitzmann, Hansjörg Baurecht

**Author notes:** Corresponding author: Sebastian E. Baumeister, PhD, Chair of Epidemiology, Ludwig-Maximilians-Universität München, UNIKA-T Augsburg, Neusässer Str. 47, 86156 Augsburg, Germany. contributed equally (Michael F Leitzmann, Hansjörg Baurecht).

## Abstract

**Introduction:** Evidence from observational studies for the effect of physical activity on the risk of Alzheimer’s disease (AD) is inconclusive. We performed Mendelian randomization analysis to examine whether physical activity is a protective factor for AD.

**Methods:** Summary data of genome-wide association studies on physical activity and AD were identified using PubMed and the GWAS catalog. The study population included 21,982 AD cases and 41,944 cognitively normal controls. Eight single nucleotide polymorphisms (SNP) known at P < 5×10^−8^ to be associated with accelerometer-assessed physical activity served as instrumental variables.

**Results:** Genetically predicted accelerometer-assessed physical activity had no effect on the risk of AD (inverse variance weighted odds ratio [OR] per standard deviation (SD) increment: 1.03, 95% confidence interval: 0.97-1.10, *P*=0.332).

**Discussion:** The present study does not support a relationship between physical activity and risk of AD, and suggests that previous observational studies might have been biased.

## Introduction

Alzheimer’s disease (AD) is the main cause of dementia and one of the great health-care challenges of the 21^st^ century [1]. Research since the discoveries of amyloid-β and tau, the main components of plaques and tangles, has provided considerable knowledge about molecular pathways of AD development; however, this knowledge has not yet been translated into the implementation of effective prevention measures for modifiable risk factors of AD [2]. Considerable research has focused on the potentially protective role of physical activity for AD. Several meta-analyses of observational studies suggested a protective effect of physical activity for cognitive decline and risk of dementia and AD [3-10]. Although intervention studies have shown that exercise improves cognitive performance, they have not revealed changes in the risk of dementia or AD [4, 11]. For example, the large multidomain lifestyle FINGER (Finish Geriatric Intervention Study to Prevent Cognitive Impairment and Disability) trial comprised an exercise program and demonstrated beneficial effects on cognition after two years [12].

More recently, long-term observational studies have suggested that the inverse association of physical activity and dementia might be subject to reverse causation due to a decline in physical activity during the preclinical phase of dementia [13, 14]. Mendelian randomization (MR) is a method that uses genetic variants as instrumental variables to uncover causal relationships in the presence of observational study bias such as unobserved confounding and reverse causation [15]. In the current study, we performed two-sample MR analyses to provide further evidence for the association between accelerometer-assessed physical activity and AD.

## Methods

The MR study design had three components: (1) identification of genetic variants to serve as instrumental variables for accelerometer-assessed physical activity; (2) the acquisition of summary data for the genetic instruments from genome-wide association studies on accelerometer-assessed physical activity; (3) acquisition of instrumenting SNP-outcome summary data for the effect of genetic instruments from genome wide association studies on the risk of AD.

### Instrumental variables for accelerometer-assessed physical activity

We selected eight SNPs associated with accelerometer-based physical activity (mean acceleration in milli-gravities) at a genome-wide significance level (P < 5 × 10^−8^), using a PLINK-clumping algorithm (r^2^ threshold = 0.001 and window size = 10mB), from a genome-wide study of 91,084 UK Biobank participants [16] (Supplementary Table 1).

### GWAS summary data for AD

Summary data for the association of SNPs for accelerometer-based physical activity with AD were obtained from a GWAS of 21,982 clinically-confirmed AD cases and 41,944 cognitively normal controls [17]. That GWAS for AD did not include the data from the UK Biobank.

## Statistical power

The a priori statistical power was calculated using an online tool at http://cnsgenomics.com/shiny/mRnd/ [18]. We assumed that the eight accelerometer-based physical activity SNPs explained 0.4% of the phenotypic variable [16, 19, 20]. Given a type 1 error of 5%, we had sufficient statistical power (>85%) for an expected odds ratios (OR) per 1 standard deviation of ≤0.88 between AD and genetically instrumented accelerometer-based physical activity.

## Statistical analyses

Cochran’s Q was computed to quantify heterogeneity across the individual effect estimates of the selected SNPs, with P≤ 0.05 indicating the presence of pleiotropy (Supplemental Table 2). Consequently, a random effects inverse-variance weighted (IVW) MR analysis was used as principal analysis [21]. Other MR methods addressing specific violations of specific instrumental variable analysis assumptions included: weighted median, MR-Egger and MR-Pleiotropy RESidual Sum and Outlier (MR-PRESSO) [22, 23]. The results were presented as ORs and 95% confidence intervals (CIs) per 1-SD increment in accelerometer-based physical activity. We tested for potential directional pleiotropy by testing the intercepts of MR-Egger models [22]. Finally, we looked up each instrument SNP and its proxies (r^2^>0.8) in Phenoscanncer [24] and the GWAS catalog [25] to assess any previous associations (P<1×10^−8^) with potential confounders. We performed leave-one-out analyses and exclusion of potentially pleiotropic SNPs to rule out possible pleiotropic effects. Analyses were performed using the TwoSampleMR (version 0.4.25) [23] and MRPRESSO (version 1.0) packages in R (version 3.6.1). Reporting follows the STROBE-MR statement [26].

## Results

We found that genetically predicted accelerometer-based physical activity was not associated with AD (IVW OR per 1-SD increment: 1.03, 95% CI: 0.97-1.09 *P*=0.334, Table 1). This finding was confirmed using alternative MR methods and leave-one-SNP-out-analysis (Table 1, Supplementary Table 3). The F-statistics for the strength of the genetic instruments were all ≥10 and ranged from 30 to 48 (Supplementary Table 1). The intercept test from the MR-Egger regression was not statistically significant (Supplementary Table 2). In the Phenoscanner and GWAS databases, we did not find an indication of possible pleiotropy of any of the eight SNPs for accelerometer-based physical activity.

**Table 1.** Mendelian randomization estimates between accelerometer-based physical activity and Alzheimer’s disease

|  | Method | OR <sup>a</sup> | 95% CI | P value |
| --- | --- | --- | --- | --- |
| Alzheimer's disease |  |  |  |  |
|  | Inverse-variance weighted (random effects) | 1.03 | 0.97; 1.10 | 0.332 |
|  | Weighted median | 1.05 | 0.98; 1.12 | 0.189 |
|  | MR-Egger | 1.11 | 0.83; 1.49 | 0.513 |
|  | MR-PRESSO | 1.03 | 0.97; 1.10 | 0.363 |

OR (odds ratio) per one standard deviation increment in mean acceleration (in milli-gravities). CI, confidence interval.

## Discussion

This MR study with GWAS data on accelerometer-based physical activity from 91,084 individuals and GWAS data from 21,982 AD cases found little evidence for an effect of physical activity on the risk of developing AD. Previous observational studies concluded that higher levels of physical activity are associated with reduced risk of dementia and AD [6, 7, 9]. The most comprehensive meta-analysis comprising 15 cohort studies found a 35% (relative risk: 0.65; 95% CI: 0.56-0.74) relative reduction in risk of AD when comparing the highest and lowest levels of physical activity [7]. These conclusions are in contrast to meta-analyses of intervention studies, which do not show a protective effect of exercise interventions on the risk of AD [4, 11]. Similarly, recent observational studies have found that when physical activity assessment and diagnosis of AD are ≥10 years apart there was no association between physical activity and risk of dementia and AD [13, 14]. A pooled analysis [13] of 19 studies including 1,602 AD cases found a hazard ratio of 1.04 (95 CI: 0.91; 1.19) when comparing physically active and inactive individuals and restricting follow-up time to ≥10 years. Furthermore, the latter studies also have indicated that a decline in physical activity levels occurs during the subclinical or prodromal phase of dementia and that previous observational studies might have overestimated dementia risk associated with insufficient levels of physical activity as many studies were based on short follow-up times and thus may have been subject to reverse causation caused by decline in physical activity prior to the diagnosis of dementia [13]. We conducted an MR analysis, which is less susceptible to reverse causation, to further shed light on the association between physical activity and AD. The findings of the present study do not suggest a causal effect of physical activity on AD.

Our study had several strengths. The use of two-sample MR enabled us to use the largest GWAS on AD to date. Our MR study also incorporated the largest GWAS on physical activity to increase the precision of SNP-physical activity estimates, to reduce the potential for weak instrument bias and to increase statistical power. We used genetically predicted objectively measured physical activity, using accelerometry, which is less prone to recall and response bias than measurement of self-reported physical activity [27]. Furthermore, because some genetic loci for self-reported physical activity are also related to cognitive function, self-reported physical activity measures may be prone to information bias, and SNP instrumenting self-reported physical activity might have induced horizontal pleiotropy [16, 28]. In contrast, SNP-associations based on accelerometer-assessed physical activity are unrelated to cognitive performance or other potential pathways with AD, which essentially rules out any impact cognitive biases or pleiotropy could have had on our results [16, 28]. However, our study also had certain limitations. First, the genetic instruments for accelerometer-assessed physical activity explained only a small fraction of the phenotypic variability. Second, for the two-sample MR to provide unbiased estimates, the risk factor and outcome sample should come from the same underlying population. The discovery genome-wide association study of physical activity consisted of UK Biobank participants of European descent, aged 40 to 70 years [16]. The SNP-AD associations were derived from cohort and case-control studies of men and women of European descent aged 65 years and older [17]. By using non-specific effects, our analyses assumed that the effects of SNPs on physical activity do not vary by age. However, this may not be an entirely tenable assumption given that the heritability of physical activity has been shown to decrease with age [29]. Thus, given the limited age range of the UK Biobank and inclusion of European ancestry individuals only, our results may not be generalizable to other age groups or ancestral populations. Therefore, replication of our findings in other age groups and non-European populations is warranted.

Given the increase in life expectancy, AD is increasingly a public health challenge and measures to prevent or delay the onset of dementia are urgently needed. However, in combination with previous literature [13, 14], the present study provides little evidence that recommending physical activity would help to prevent AD.

## Conflicts of Interest Disclosures

All authors disclose no conflict.

## Funding/Support

The authors did not receive funding for this study. Funding information of the genome-wide association studies is specified in the cited studies.

## Data availability

Data supporting the findings of this study are available within the paper and its supplementary information files.

